# MicroRNA-374b regulates SARS-CoV-2 viral protein mediated endothelial to mesenchymal transition by targeting c-FLIP

**DOI:** 10.1101/2024.05.22.595176

**Authors:** Grace R. Raji, Aswini Poyyakkara, Vishnu Ramachandran, K Anjali, VB Sameer Kumar

## Abstract

The pathophysiological consequences of COVID-19 disease are still unclear, however, endothelial cell (EC) dysfunction has been observed to play a key role in disease progression and severity. Many reports suggests that SARS-CoV-2 mediated endothelial dysfunction is the result of intracellular signaling initiated by the binding of the spike protein to ACE2, which can modify endothelial cell phenotype. Recent reports suggests endothelial to mesenchymal transition (Endo MT) as a process heavily involved in lung fibrosis of COVID 19 patients. EndoMT is involved in many chronic and fibrotic diseases and appears to be regulated by complex molecular mechanisms and different signaling pathways, in particular microRNAs (miRNAs), which constitute a crucial mediator of EndoMT. MicroRNAs (miRNAs), small endogenous RNA molecules that regulate several physiological processes including endothelial homeostasis, and vascular diseases, can be perturbed by infecting viruses. Based on these facts, this study was designed to decipher the role of miR-374b, which was found to be significantly downregulated upon profiling of SARS-CoV-2 viral protein stimulated endothelial cells. Gene profiling of endothelial cells revealed c-FLIP (CFLAR) to be among the most significantly upregulated gene. In silico target prediction analysis using targetscan revealed c-FLIP as the major target of miR-374b. Further it was identified that miR-374b can reverse c-FLIP mRNA and protein levels in SARS-CoV-2 viral protein stimulated endothelial cells under conditions of miR-374b overexpression. Since vascular dysfunction involve, under many circumstances, loss of vascular tone due to mesenchymal transition of endothelial cells, we next checked if fibrotic events are initiated downstream of c-FLIP pathway. Further mechanistic studies involving identification of the expression pattern of mesenchymal markers in SARS-CoV-2 viral protein stimulated endothelial cells in presence or absence of miR-374b provide evidence for the important role of miR-374b in regulating SARS CoV-2 mediated EndoMT and fibrotic events downstream of c-FLIP pathway and may highlight possible new therapeutic approaches targeted at the damaged endothelium.

## 1. Introduction

Coronavirus disease 2019 (COVID-19) represents a public health crisis caused by the severe acute respiratory syndrome coronavirus 2 (SARS-CoV-2). SARS-CoV-2 is a positive single stranded RNA virus with large genome of 29.9kb causing acute respiratory distress syndrome (ARDS) which is the most common cause for high mortality rates [1]. Although SARS-CoV-2 primarily affects the pulmonary system, growing evidences suggests that it can also impact pan-vasculature and contribute to a number of extrapulmonary manifestations, including macro and micro-thromboembolic events [2] and disseminated intravascular coagulation in patients with COVID-19 leading to multi-organ failure [3]. Recent reports suggests systemic involovement of COVID-19 can be related to endothelial dysfunction (endothelitis, endothelialitis and endotheliopathy)[4] caused by either direct viral effects or virus-dependent induction of the inflammatory response [5].

SARS-CoV-2 mediated endothelial dysfunction is the result of intracellular signaling initiated by the binding of the spike protein to ACE2 [6], which can modify endothelial cell phenotype, thus leading to multiple instances of endothelial dysfunction, including oxidative stress, reduced nitric oxide bioavailability, glycocalyx/barrier disruption, hyperpermeability, inflammation/leukocyte adhesion, hypercoagulability, thrombosis and endothelial-to-mesenchymal transition [7][8]. Mounting evidences suggests endothelial to mesenchymal transition (Endo MT) as a process heavily involved in lung fibrosis of COVID 19 patients [9]. EndoMT is a cellular differentiation process in which endothelial cells (ECs) lose their endothelial specific properties and acquire mesenchymal properties, thereby showing decreased expression of endothelial-specific cell adhesion markers, such as vascular endothelial (VE)-cadherin and increased expression of mesenchymal markers like N-cadherin, Vimentin and SMA, thus enhancing the migratory property of the cells [10]. EndoMT is reported to play a key role in tissue stiffening by expressing extracellular membrane (ECM) proteins. During inflammation, various factors such as inflammatory cytokines, reactive oxygen species etc, continuously activates ECs leading to altered vascular relaxation, increased leucocyte adhesion and endothelial permeability resulting in a prothrombotic state, ultimately leading to endothelial dysfunction[11][12]. EndMT appears to be regulated by complex molecular mechanisms and different signaling pathways, including microRNAs (miRNAs)[13].

MicroRNAs (miRNAs), small endogenous RNA molecules that regulate several physiological processes including endothelial homeostasis, and vascular diseases, can be perturbed by infecting viruses [14]. It has been reported that, SARS-CoV-2 infection can alter host miRNA expression landscape, resulting in miRNA mediated regulation of target mRNA and its downstream signaling cascade [15]. Eventhough host miRNAs are reported to be associated with SARS-CoV-2 infection, role of host miRNAs in regulating SARS-CoV-2 mediated endothelial dysfunction still remains elusive. Based on these facts, this study was designed to decipher the role of miRNAs in SARS-CoV-2 viral protein stimulated endothelial cells. In particular, this study focus on the role of miR-374b, which was found to be significantly downregulated upon profiling of SARS-CoV-2 viral protein, stimulated endothelial cells. Eventhough, miR-374b play crucial role in Endo MT and also has been reported to be downregulated in blood samples of COVID 19 patients [16], the role of miR-374b in SARS-CoV-2 mediated endothelial dysfunction remains unclear and this forms the subject matter of the present study.

## 2. Materials and methods

### Materials

Human umbilical vein endothelial cell lines (HUVEC/TERT2) was procured from ATCC. HEK293T cells were obtained from NCCS, Pune. SARS-CoV2 spike (https://www.addgene.org/145780) and SARS-CoV2 nucleocapsid (https://www.addgene.org/158079) expressing plasmids were purchased from Addgene. pCMVMIR vector was obtained from OriGene, USA. HiEndoXL Endothelial cell expansion medium, endothelial growth supplement, DMEM (Dulbecco’s Modified Eagles Medium), Opti MEM, FBS (Fetal Bovine Serum), BSA (Bovine Serum Albumin), Antibiotic antimycotic solution, Trypsin and Molecular biology grade water were obtained from Himedia, India. Proteinase inhibitor cocktail, Anti β-Actin antibody and c-FLIP rabbit polyclonal antibody (PROMPT) was obtained from origin diagnostics, Kerala. Anti Mouse IgG, Tween-20, Acrylamide, SDS (Sodium dodecyl sulphate), N’ methylene bis-acrylamide, ammonium persulphate, TEMED (N, N, N’, N’-Tetra methyl ethylene diamine), actinomycin, were purchased from Sigma Aldrich Co. USA. Human miRnome miScript miRNA PCR array, miRNA isolation kit, miScript II RT kit, QuantiTect SYBR GREEN PCR master mix were procured from Qiagen, Germany and cDNA Reverse transcription kit was procured from Biorad. Lipofectmine reagent and Drabkins reagent, propidium iodide were obtained from Invitrogen. All the solutions and reagents were prepared using ultrapure Milli-Q water.

### Methods

#### Cell culture

Human Umbilical Vein Endothelial cells (HUVECs) were cultured in HiEndoXL Endothelial cell expansion medium containing Endothelial Growth Supplement and 1% Antibiotic Antimycotic solution. HEK293 cell line was cultured in DMEM supplemented with 10% FBS and 1% antibiotic antimycotic solution and L-glutamine. The cells were maintained under standard culture conditions of humidified atmosphere with 95% air and 5% CO_2_ at 37°C. The experiments were performed by subjecting the cells to different treatment conditions.

#### Profiling of host miRNAs from SARS-CoV-2 viral protein stimulated endothelial cells

Human umbilical vein endothelial cells (HUVECs) cultured in 6 well plates under standard conditions were co-transfected with SARS-CoV2 N plasmid and pEGFP-C1 using lipofectamine LTX reagent as per manufacturer’s instructions. 6 hours post transfection, media was changed and replaced with fresh media containing spike protein (200ug). To determine the efficiency of transfection, cells were observed for green fluorescence of pEGFP-C1 under a fluorescent microscope. 48 hours post transfection, cells were harvested and miRNAs were isolated using miRNA isolation kit (Qiagen) as per manufacturer’s instruction. The concentration of RNA was measured using nanophotometer. 1ug of small RNA was used for cDNA preparation using miscript II RT kit (Qiagen) as per the manufacturer’s instructions and profiled the most abundant 1092 miRNAs using human mirnome miscript miRNA PCR array (Qiagen). The data was analysed by ΔΔCT method of relative quantification using an analysis tool available at geneglobe.qiagen.com.

#### Target prediction, 3’UTR analysis and mRNA stability assay

Target prediction of miR-374b was determined using 3 different web based tools like miRDB (mirdb.org), Targetscan (www.targetscan.org) and miRsystem (mirsystem.cgm.ntu.edu). 3’UTR analysis of the identified candidate target was performed using online web based tool targetscan. For mRNA stability assay, HEK293 cells were seeded at a density of 2 x 10^5^ cells/well in 6 well plates. The cells after reaching a confluency of 70 % were transfected with miRNA expressing constructs (miR-374b) along with proper control vectors (pcmvmiR) using PEI method. 24 hours post transfection, cells were treated with actinomycin (6μg) for different time intervals like, 0, 1hr, 3hr, 6hr and 12hr to inhibit transcription followed by RNA isolation and RT-PCR analysis of target mRNA.

#### RNA isolation and RT-PCR analysis

Total RNA from cells cultured under different treatment conditions were isolated using TRIzol method and quantified by Nanodrop spectrophotometer (Thermo scientific). 1μg of total RNA was used for cDNA synthesis using cDNA synthesis kit. The cDNA was then diluted in a ratio of 1:10 followed by Real-Time PCR in Roche light cycler 480 using SYBR Green chemistry. Appropriate internal control and non-template controls were included in all experiments. The primers used were custom designed.

#### Western blotting analysis

Cells grown under different treatment conditions were harvested and lysed in RIPA buffer and then sonicated. The cell lysate was then collected, SDS PAGE loading dye was added and then heated at 90 °C for 10 min. Protein normalized samples were then separated on 10% SDS PAGE and then transferred on to PVDF membrane in a Trans blot apparatus. The membrane was then blocked with 5% BSA for 1 hour at room temperature and incubated with primary antibody overnight at 4°C. Further, the membranes were washed with TBST and incubated with HRP conjugated secondary antibody for 1 h. The blots were then developed using chemiluminescent reagent, followed by visualization and quantification of bands in a Versa Doc imaging system.

#### Comet assay

DNA fragmentation associated with apoptosis was determined by comet assay. HUVECs, at a density of 10^5^cells/ml, were mixed with 1% low melting agarose at a ratio of 1:3 and evenly spread over a glass slide and then allowed to become gel. After gelling, slides were placed in lysis solution (2% SDS, 0.5M Na_2_EDTA, 0.5mg/ml proteinase K at pH-8) and incubated at 37°C overnight. After overnight incubation, slides were washed in TAE buffer, and electrophoresed for 20 minutes at 12v and 400mA. Slides were then stained with propidium iodide solution (20 mg/ml) for 20 minutes and observed under fluorescent microscope. The image analysis and quantification of the DNA damage in terms of comet length, tail length, tail DNA % was performed using open comet software available at www.cometbio.org.

#### Exosome isolation from culture media

Exosomes were isolated from culture media using total exosome isolation reagent according to manufacturer’s instructions. Briefly, cell culture media was collected and centrifuged at 200g for 30min to remove the cell debris. The supernatant thus obtained was mixed with half volume of total exosome isolation reagent, and incubated 4°C at overnight. After overnight incubation, samples were centrifuged at 10,000g for 1hour at 4°C. The supernatant was removed and pelleted exosomes were then resuspended in 1X PBS, followed by protein estimation.

#### Chick chorioallantoic membrane (CAM) assay

CAM assay was performed using fertilized chick eggs, for which the fertilized eggs were incubated in an incubator at 37°C with a relative humidity of 80%. On the 10^th^ day of embryonic life (after development of complete vasculature), eggs were opened on the air sac side, removed the shell with forceps, to treat the CAMs with spike protein and exosomes, followed by incubating the egg for 2 days. The eggs were then opened on 12^th^ day of embryonic life, and the CAMs were photographed. The level of hemoglobin in the CAM was estimated using Drabkin’s reagent as a measure of vessel density and was normalized to the protein content in the CAM. Vascular density was quantified using the software AngioTool64 and the expression levels of EndoMT markers such as SMA and FSP were assessed by RT-PCR analysis.

### Statistical analysis

Results are expressed as mean with standard error of mean. The statistical significance of difference was calculated by Duncan’s One-Way Analysis of Variance (ANOVA) using the SPSS 11.0 Software. A value of *P* < 0.05 was considered significant.

## 3. Results

### Profiling of miRNAs from SARS-CoV-2 viral protein stimulated endothelial cells

In order to identify the miRNAs that are differentially expressed by SARS-CoV-2 viral protein stimulated endothelial cells and control endothelial cells, miRNAs were isolated from SARS-CoV-2 N overexpressing cells treated with SARS-CoV-2 spike protein, followed by cDNA synthesis and profiling of 1092 miRNAs using human mirnome miscript miRNA PCR array. Profiling results revealed that, among the 1092 miRNAs profiled, 800 miRNAs were found to be upregulated and 4% of these miRNAs showed greater than 100 fold upregulation in their levels. 250 miRNAs, on the other hand were found to be downregulated and 94% of these miRNAs exhibited a fold change of > -100. (Fig.1). Out of these miRNAs, miRNAs which exhibited a fold change of >100,000 include miR-106b-5p, miR-301b-3p, miR-95-3p, miR-26b-5p and let-7a-5p and fold change of > -100,000 include miR-374b, miR-1281, miR-1285, miR-1287, miR-93-3p, miR-373-5p, miR-205-3p and miR-1203. Among the significantly downregulated miRNAs, miR-374b was selected as the candidate miRNA for further studies, since it is also reported to be downregulated in blood samples of COVID-19 patients and well reported in endothelial dysfunction and thrombosis. The whole profiling data is given in the supplementary material (Table S1).

**Fig. 1:**
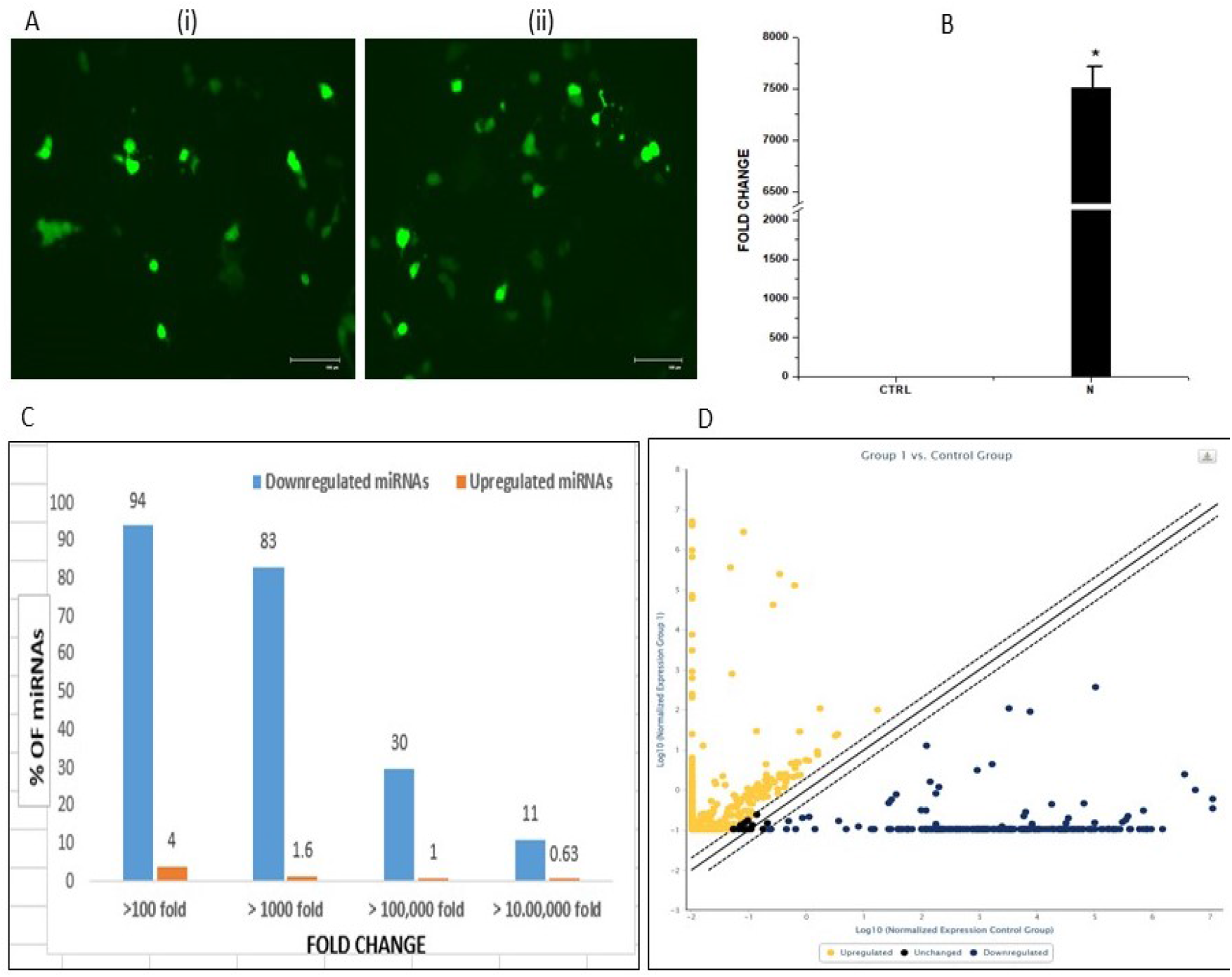
Profiling of miRNAs from endothelial cells overexpressing SARS-CoV-2 N protein and treated with SARS-CoV-2 spike protein. miRNAs were isolated from endothelial cells overexpressing SARS-CoV-2 N protein and treated with SARS-CoV-2 spike protein, converted to cDNA and profiled to check the levels of 1092 miRNAs using human mirnome miscript miRNA PCR array and data was analysed by ΔΔCT method of relative quantification using a software available at geneglobe.qiagen.com. (A) Microphotographs of (i). mock transfected cells (ii). SARS-CoV-2-N transfected cells (B) RT-PCR analysis of SARS-CoV-2-N (C) Percentage of miRNAs exhibiting higher or lower levels in endothelial cells overexpressing SARS-CoV-2 N protein and treated with SARS-CoV-2 spike protein and their fold change (D) Scatter plot analysis of miRNA profiling for differentially expressed miRNAs. miRNA profiling data after normalisation with housekeeping genes, the final scores for each of the miRNA in the test array were compared with that of the control array. The y-axis represents log scores of Test (Group 1) and x-axis represents log scores of control (Group 2). Each symbol represents individual miRNA. Those miRNAs outside the boundaries represent 2-fold higher or lower expression in test group.

### Profiling of genes associated with endothelial function and KEGG pathway analysis

In order to check, whether mimicking of viral stimulation in endothelial cells by SARS-CoV-2 S and N protein capable of modulating the expression pattern of genes associated with endothelial function and thrombosis, RT-PCR analysis of 43 genes implicated in endothelial function and thrombosis was carried out. Among the 43 genes profiled, out of the upregulated genes, HIF1A, KDR, MMP9, APOE, ACE, C-FLIP (CFLAR), TNFSF10 was found to be most significantly upregulated (> 5 fold) and out of the downregulated genes, VEGFA, THBS1, BCL2L, PECAM1, was found to be most significantly downregulated (Fig. 2A). Results suggests that most of the genes were significantly dysregulated indicating that SARS-CoV-2 S or N protein mediated endothelial dysfunction may incvolve modulation of the expression levels of multiple genes.

**Fig. 2.**
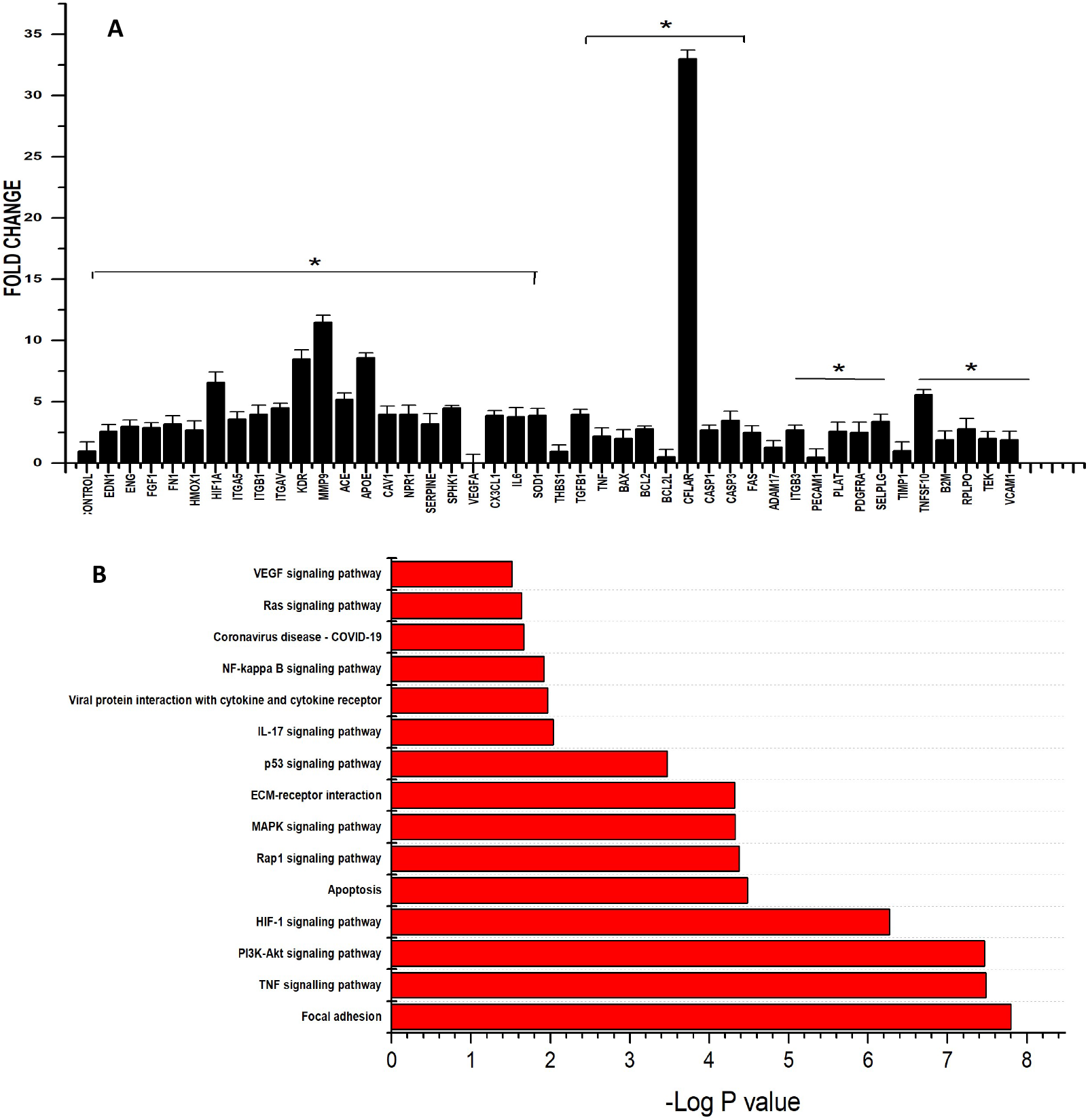
Profiling of genes associated with endothelial function and thrombosis. **(A)** RT-PCR analysis of 43 genes associated with endothelial function and thrombosis. (B) KEGG pathway enrichment analysis. 15 pathway categories (p<0.05) were significantly affected by candidate genes and 4 pathway categories were most significantly enriched. The y-axis represents –log p-value and x-axis represents different pathways.

KEGG Pathway analysis of significantly dysregulated genes performed using DAVID tool revealed that, a total of 70 pathways (p<0.1) were associated with these candidate genes. Among these 70 pathways, 15 pathway categories associated with cell proliferation or apoptosis and covid-19 disease were enriched with a p-value of <0.05 (Fig. 2B). However 4 pathway categories like focal adhesion, PI3-AKT signalling pathway, TNF signalling pathway and HIF1A signaling pathway were found to be the most significantly enriched. These pathways have also been reported to be associated with endothelial dysfunction and thrombosis.

### Expression level of miR-374b in SARS-CoV-2 viral protein stimulated ECs: Prediction of its target

Subsequent to our observation that, miR-374b was significantly downregulated upon profiling and was reported to be downregulated in blood samples of COVID-19 patients, we selected miR-374b as candidate miRNA and verified the expression levels of miR-374b in SARS-CoV-2 viral protein stimulated ECs. The results reaffirmed significant downregulation in the expression level of miR-374b in SARS-CoV-2 viral protein stimulated ECs (Fig. 3A). Further we, predicted the targets of miR-374b, using 3 online web based tools like miRDB, targetscan and miRsystem, followed by scoring of the identified targets. Among the predicted targets, only the genes associated with endothelial function and involved in gene profile were selected for further analysis. Target prediction analysis revealed CFLAR (c-FLIP), ITGB1 and VEGFA as the important targets. The target gene identified by all the three independent tools was selected for further mechanistic study. Among the candidate targets, only CFLAR(c-FLIP) could satisfy our criteria with a score of 3/3 (Fig.3B). Also among the candidate genes, CFLAR was also found to be the most significantly upregulated gene in gene profiling, and therefore was selected for further mechanistic studies. CFLAR (c-FLIP) being predicted as the target of miR-374b, we next checked c-FLIP mRNA and protein levels in SARSCoV2 viral protein stimulated ECs. For this SARS-CoV2 N transfected ECs were treated with SARSCOV2 spike protein, followed by analysis of CFLAR mRNA levels by RT-PCR and protein levels by western blotting. Results obtained showed that c-FLIP mRNA and protein levels were significantly inceased in SARSCoV-2 viral protein stimulated ECs, when compared to untreated control ECs (Fig. 3C). Further, 3′UTR analysis results revealed that 3′UTR of c-FLIP harbours two putative miR-374b binding regions at nucleotides 6160–6167 and 10988–10994, suggesting that 3′UTR of CFLAR (c-FLIP) possess potential binding sites for miR-374b (Fig. 3D). In order to check whether CFLAR is targeted by miR-374b, CFLAR mRNA stability assay was carried out, and the results obtained showed that the stability of CFLAR mRNA was significantly reduced under miR-374b overexpression condition, suggesting that the regulatory role of miR-374b involves alterations in the post-transcriptional stability of CFLAR mRNA (Fig. 3E)

**Fig. 3.**
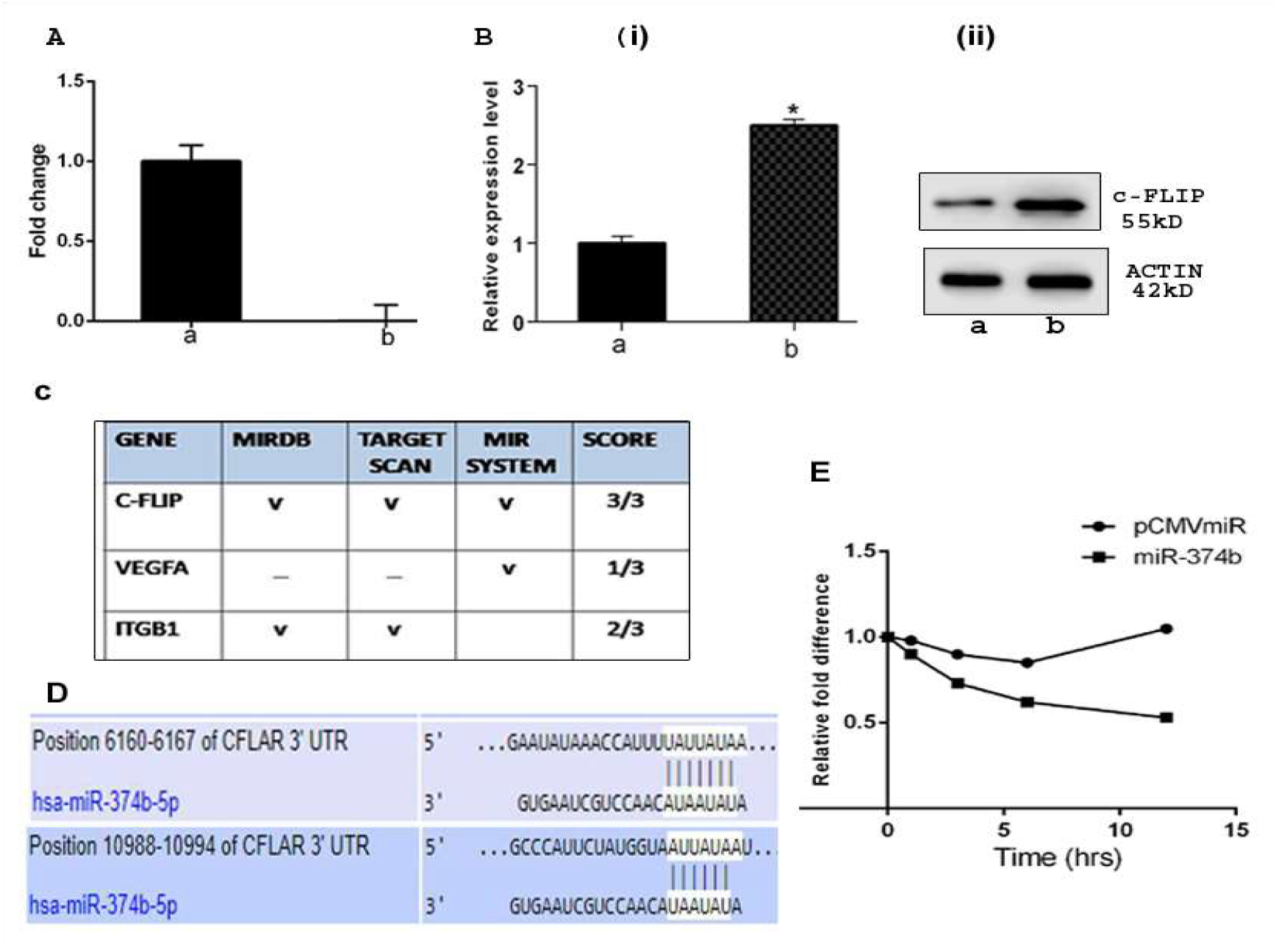
Expression levels of miR-374b, c-FLIP 3’UTR analysis and mRNA stability assay. **A.** Expression level of miR-374b in SARSCoV2 viral protein stimulated ECs. **B**. CFLAR mRNA and protein levels in SARSCoV2 viral protein stimulated ECs. For this SARS-CoV2 N transfected ECs were treated with SARSCOV2 spike protein, followed by analysis of CFLAR mRNA levels by RT-PCR and protein levels by western blotting. In A and B, a represents untreated control ECs and b represents SARS CoV2 viral protein stimulated ECs (endothelial cells overexpressed with SARS-CoV-2 N protein and treated with SARS-CoV-2 spike protein) **C**. miR-374b target prediction. Target prediction was carried out using 3 online web based tools like miRDB, targetscan and miRsystem, followed by scoring of the identified targets. **D**. 3 ‘UTR analysis of CFLAR. CFLAR 3’UTR analysis was carried out using a web based tool targetscan. This figure shows the sequence alignments between miR-374b and 3’ UTR of CFLAR at two positions. **E**. CFLAR mRNA stability assay. miR-374b overexpressing HEK293 cells were treated with actinomycin at a concentration of 6μg for different time intervals like 0, 1, 3, 6 and 12hr followed by RNA isolation from these cells and RT-PCR analysis of CFLAR mRNA. Results presented are average of three experiments ± SEM each done at least in duplicate, p < 0.05. *Statistically significant when compared to a.

### miR-374b targets c-FLIP to promote apoptosis in SARSCoV-2 viral protein stimulated ECs

c-FLIP being predicted as the target of miR-374b and the levels of c-FLIP was found to be significantly increased in SARSCoV-2 viral protein stimulated ECs, we next checked the effect of miR-374b overexpression on c-FLIP mRNA and protein levels in SARSCoV-2 viral protein stimulated ECs. For this miR-374b and SARS-CoV2 N transfected ECs were treated with SARSCOV2 spike protein. 24hrs post spike protein treatment, cells were harvested, analysis of c-FLIP mRNA and protein levels by RT-PCR and western blot respectively were carried out and the results obtained showed that c-FLIP mRNA and protein levels were significantly reduced in miR-374b overexpressing SARSCoV-2 viral protein stimulated ECs, when compared to SARSCoV-2 viral protein stimulated ECs (Fig. 4B).

**Fig. 4.**
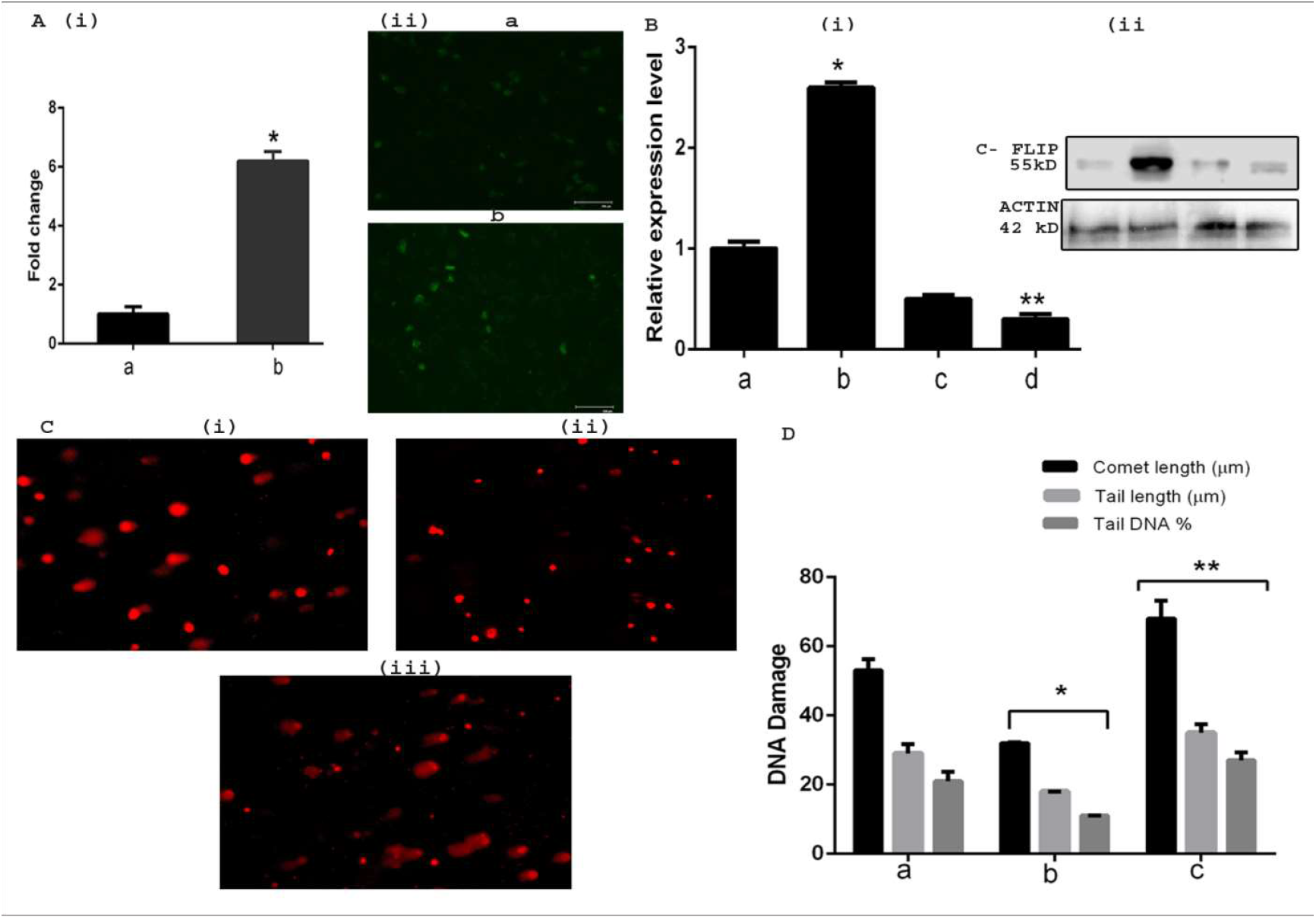
miR-374b targets c-FLIP to promote apoptosis in SARSCoV2 viral protein stimulated ECs. **A. (i)** Expression level of miR-374b in ECs.. The construct expressing miR-374b was transfected into SARSCoV2 viral protein stimulated ECs using lipofectamine reagent, followed by RT-PCR analysis of miR-374b in transfected cells. **(ii)** Microphotographs of cells transfected with **(a)** pCMVmiR (vector control) **(b)** miR-374b construct **(B)** In (A), **a** represents pCMVmiR (vector control) transfected cells **b** represents miR-374b overexpressed cells. **B**. c-FLIP mRNA and protein levels in SARSCoV2 viral protein stimulated ECs under conditions of miR-374b overexpression. For this miR-374b and SARS-CoV2 N transfected ECs were treated with SARSCOV2 spike protein. 24hrs post spike protein treatment, followed by analysis of c-FLIP mRNA levels by RT-PCR (i) and protein levels by western blot (ii). In B, a represents vector alone transfected cells, b represents vector + SARSCoV2 N + spike protein treated cells, c represents miR-374b overexpressing cells and d represents miR-374b overexpressing cells transfected with SARSCoV2 N and treated with spike protein **(C)** Representative microphotographs of comets obtained from comet assay carried out in miR-374b overexpressing SARSCoV2 viral protein stimulated ECs. **(D)** Levels of DNA damage analysed by comet assay in miR-374b overexpressing SARSCoV2 viral stimulated ECs represented in terms of different parameters like comet length,tail length and tail DNA%. The mean of the parameters were analysed from 25 independent cells per sample using open comet software. In C, a represents vector alone transfected ECs. b represents SARS-CoV2 N transfected ECs treated with SARSCOV2 spike protein (SARSCoV2 viral stimulated ECs), c represents miR-374b overexpressing SARSCoV2 viral protein stimulated ECs. Results presented are average of three experiments ± SEM each done at least in duplicate, p < 0.05. *Statistically significant when compared to a. **Statistically significant when compared to b.

c-FLIP, being an anti-apoptotic protein, we further checked the effect of miR-374b on c-FLIP mediated anti-apoptotic effect in SARSCoV-2 viral protein stimulated ECs. For this miR-374b and SARS-CoV2 N transfected ECs were treated with SARSCOV2 spike protein. 24hrs post spike protein treatment, comet assay was carried out. Results of which showed an increased rate of DNA damage or fragmentation and apoptotic levels in miR-374b overexpressing SARSCoV-2 viral protein stimulated ECs. These results clearly suggest that ectopic expression of miR-374b increased the percentage of cells undergoing apoptosis in SARSCoV-2 viral protein stimulated ECs by reducing CFLAR levels (Fig. 4D).

### miR-374b dependent regulation of EndoMT in SARSCoV-2 viral protein stimulated ECs

The results here suggest that, lower level of miR-374b and high levels of c-FLIP under conditions of SARS-CoV-2 induction is anti-apoptotic and pro-survival for the ECs suggesting its role in EC survival. Since it was also observed that, endothelial cell specifc markers like PECAM1, VWF and VE-CADHERIN was reduced in SARSCoV-2 viral protein stimulated ECs (Fig. 5A, A), Next we tried to check if the levels of miR-374b and c-FLIP could alter the endothelial function. For this RNA was isolated from SARSCoV-2 viral protein stimulated ECs, followed by cDNA synthesis and RT-PCR analysis of EndoMT markers like α SMA, Vimentin, FSP, and N-cadherin. Results of which showed that EndoMT markers were significantly increased in SARSCoV-2 viral protein stimulated ECs when compared to control cells (Fig. 5A, B). Further in order to check the role of miR-374b in modulation of EndoMT markers, we next analysed the expression pattern of EndoMT markers in miR-374b overexpressing SARSCoV-2 viral protein stimulated ECs. For this miR-374b and SARS-CoV2 N transfected ECs were treated with SARSCOV2 spike protein. 24hrs post spike protein treatment, followed by cDNA synthesis and RT-PCR analysis of EndoMT markers like α SMA, Vimentin, FSP, and N-cadherin. Results of which showed that EndoMT markers were significantly reduced in miR-374b overexpressing SARSCoV-2 viral protein stimulated ECs, suggesting that ectopic expression of miR-374b can reverse /rescue the increase in the expression pattern of EndoMT markers (Fig. 5A, C).

**Fig. 5A.**
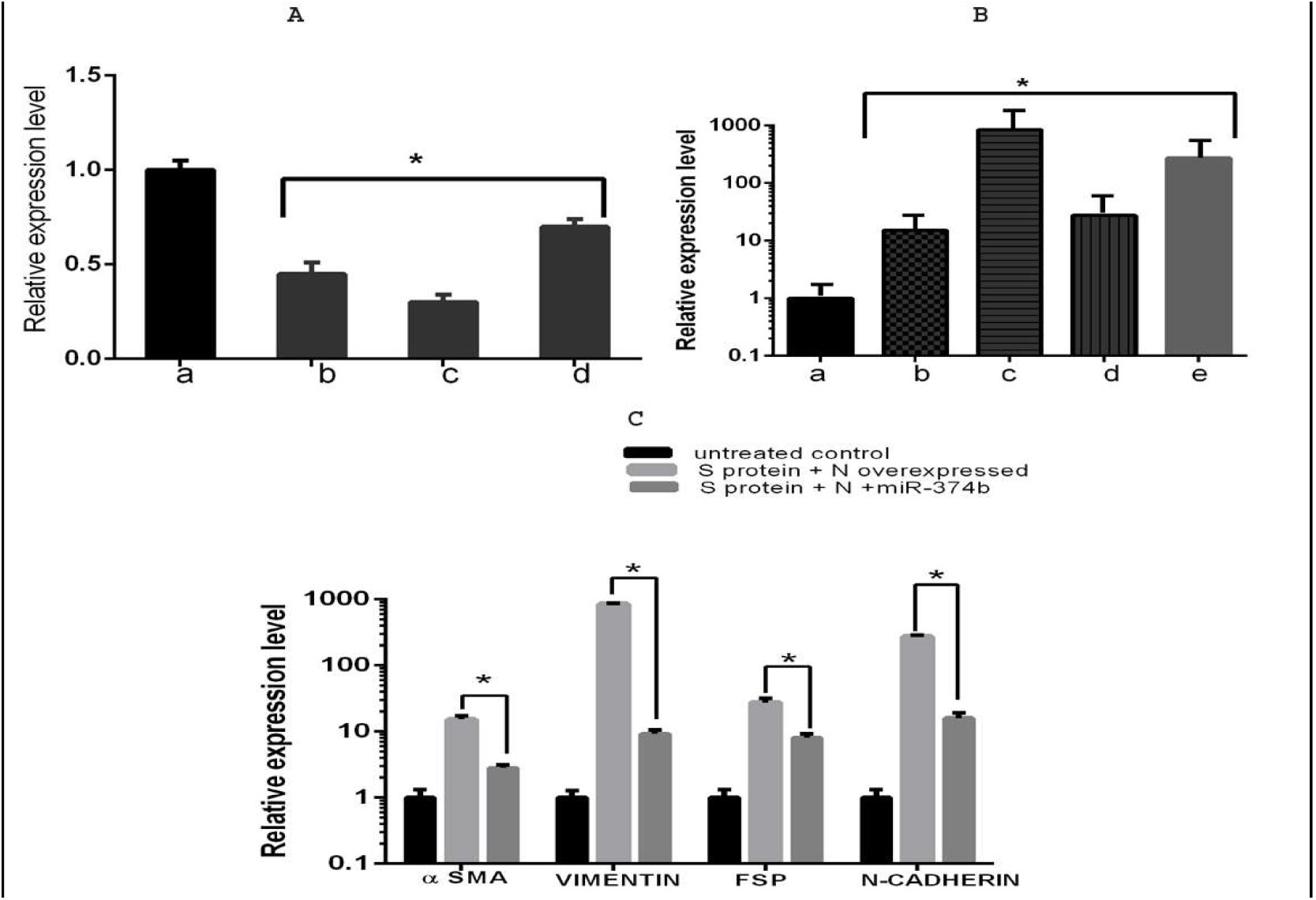
miR-374b dependent regulation of EndoMT in SARSCoV-2 viral protein stimulated ECs. **A**. Expression levels of endothelial specific markers in SARSCoV-2 viral protein stimulated ECs. For this RNA was isolated from SARSCoV-2 viral protein stimulated ECs followed by cDNA synthesis and analysis of the levels of endothelial specific markers like (b)PECAM1, (c)VWF and (d)VE-CADHERIN by RT-PCR. **B**. Expression levels of EndoMT markers in SARSCoV-2 viral protein stimulated ECs. For this RNA was isolated from SARSCoV-2 viral protein stimulated ECs followed by cDNA synthesis and analysis of the levels of EndoMT markers by RT-PCR. In B, b,c, d, and e represents SMA, VIMENTIN, FSP and N-CADHERIN respectively. **C**. Expression level of endoMT markers in SARSCoV-2 viral protein stimulated ECs under conditions of miR-374b overexpression. For this miR-374b and SARS-CoV2 N transfected ECs were treated with SARS-COV-2 spike protein. 24hrs post spike protein treatment, followed by analysis of the levels of EndoMT markers by RT-PCR. Results presented are average of three experiments ± SEM each done at least in duplicate, p < 0.05. *Statistically significant when compared to a.

**Fig. 5B.**
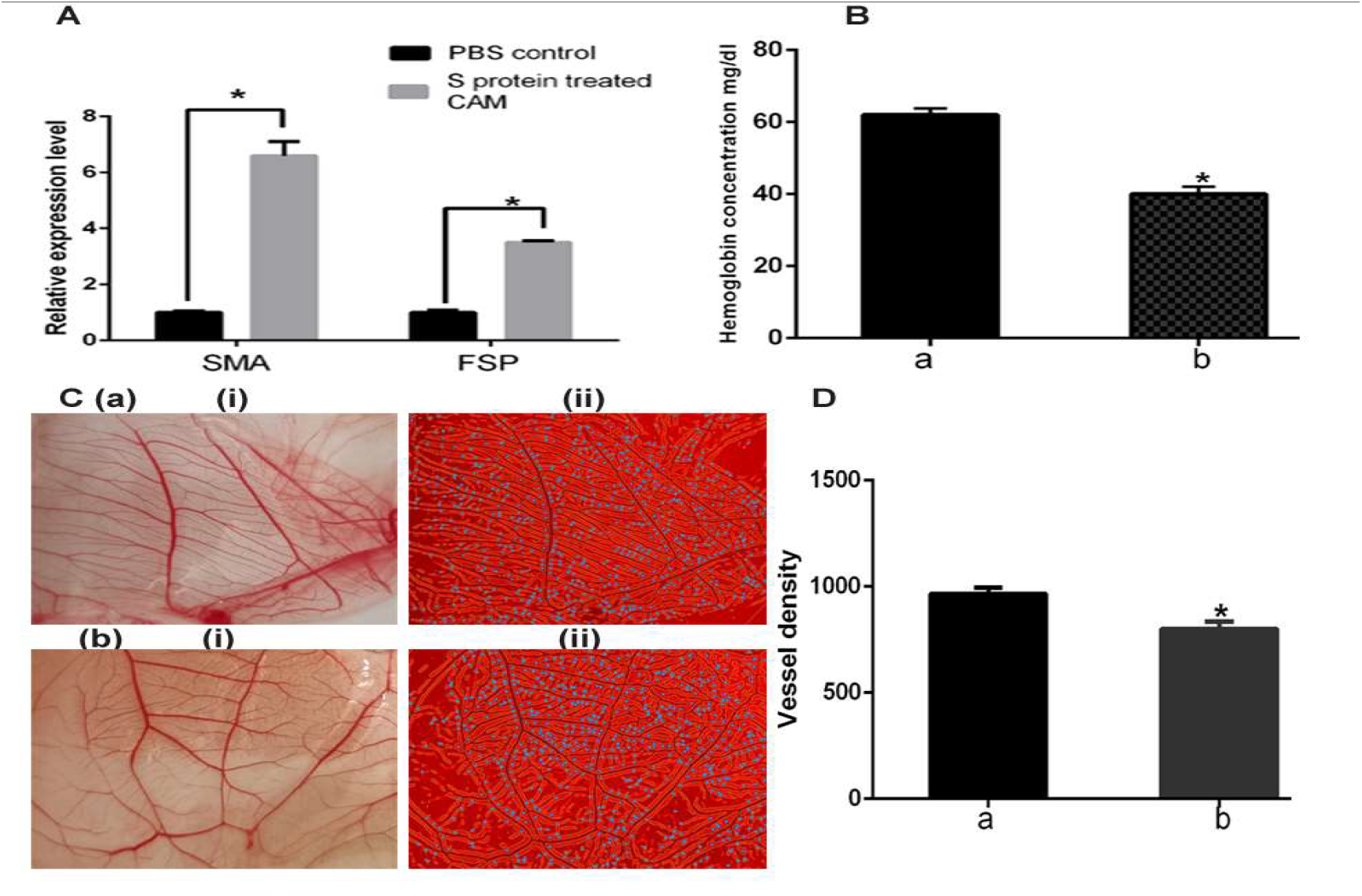
miR-374b dependent regulation of EndoMT in SARSCoV-2 viral protein stimulated ECs. **A**. Expression level of EndoMT markers in SARS CoV-2 S-protein treated CAM. For this RNA was isolated from CAM followed by cDNA synthesis and analysis of the levels of EndoMT markers like SMA and FSP by RT-PCR. **B**. Hemoglobin levels from CAM as a measure of vessel density. **C**. (i) Photographs of CAM showing vessel density (ii) Angiotool analysis of CAM showing branch points. **D**. Plot of vascular density against samples. In B, C and D, a and b represents PBS treated CAM and SARSCoV2 S-protein treated CAM respectively. Results presented are average of three experiments ± SEM each done at least in duplicate, p < 0.05. *Statistically significant when compared to a.

**Fig. 5C.**
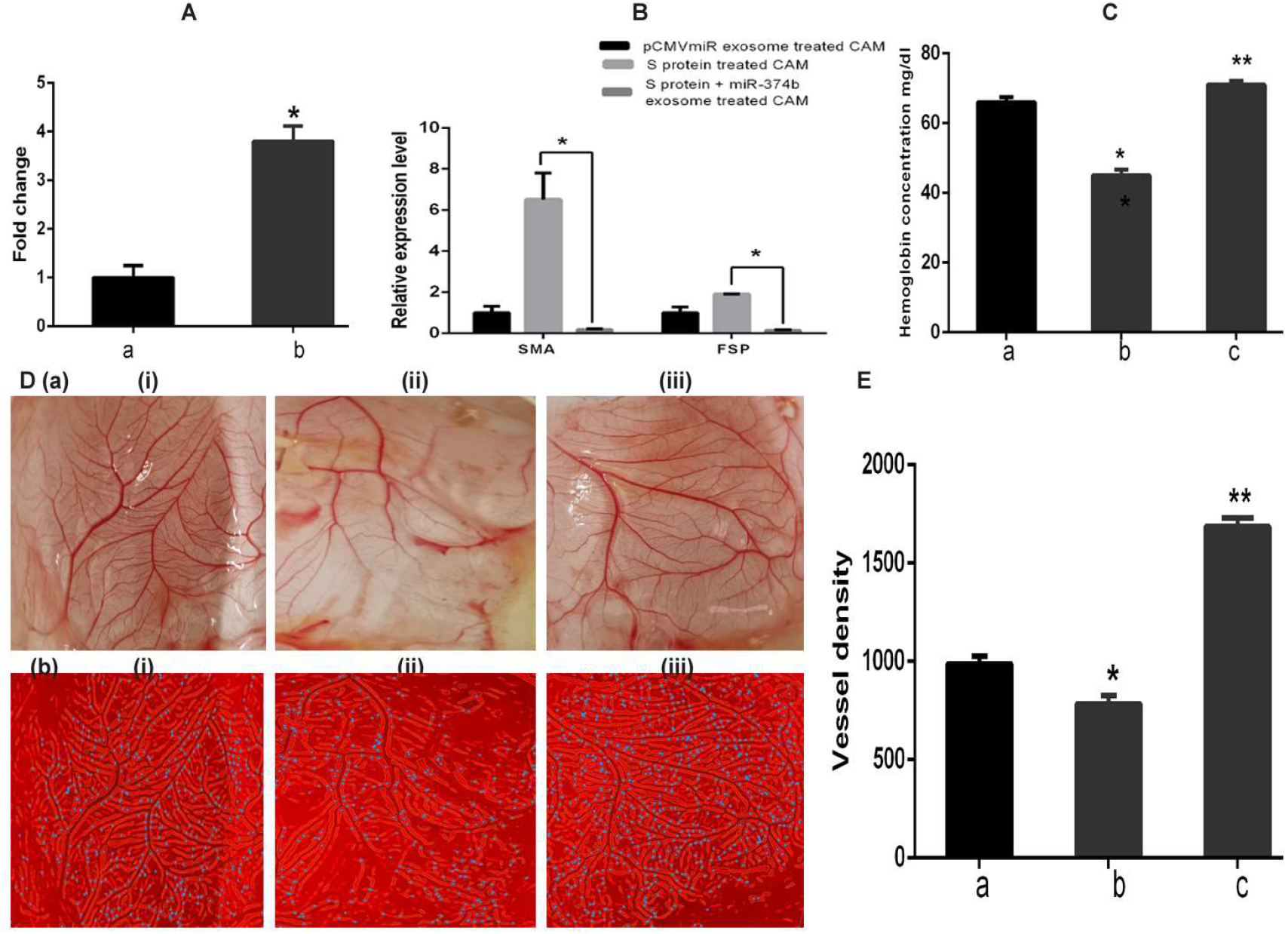
miR-374b dependent regulation of EndoMT in SARSCoV-2 viral protein stimulated ECs. **A**. Expression level of miR-374b in exosomes isolated from miR-374b overexpressing HEK293 cells. For this exosomes were isolated from the spent media of miR-374b overexpressing HEK293 cells, followed by exosomal RNA isolation, reverse transcription and and analysis of the levels of miR-374b by RT-PCR. **B**. Expression level of endo MT markers in SARS CoV-2 S-protein and miR-374b exosomes treated CAM. For this RNA was isolated from SARS CoV-2 S-protein and miR-374b exosomes treated CAM followed by cDNA synthesis and analysis of the levels of EndoMT markers like SMA and FSP by RT-PCR. **C**. Hemoglobin levels from SARS CoV-2 S-protein and miR-374b exosomes treated CAM as a measure of vessel density. **D**. (a) Photographs of CAM showing vessel density (b) Angiotool analysis of CAM showing branch points. **E**. Plot of vascular density against samples. In C and D and E, a, b and c represents pCMVmiR exosome (Exovector control) treated CAM, SARS CoV-2 S-protein treated CAM and SARSCoV2 S-protein and miR-374B exosome (EXO miR-374b) treated CAM respectively. Results presented are average of three experiments ± SEM each done at least in duplicate, p < 0.05. *Statistically significant when compared to a. **Statistically significant when compared to b

In order to further confirm the role of miR-374b in regulating Endo MT, chick Chorioallantoic membrane (CAM) assay was performed. For this CAMs of 10-day-old chick embryos were co-treated with SARSCoV2 S protein and miR-374b exosomes (EXOmiR-374b) and incubated till 12th day of embryonic life, after which the CAMs were exposed, photographed and levels of hemoglobin (as a measure of vessel density) was estimated and the expression levels of Endo MT markers (SMA and FSP) were analyzed by RT-PCR. Vascular density in the CAMs was also quantified using the software AngioTool64. Results of which showed that the expression level of Endo MT markers (SMA and FSP) increased in CAMs treated with SARSCoV2 spike protein was found to be reversed in CAMs co-treated with miR-374b exosomes. The micro vessel density and hemoglobin levels were found to be significantly lower in CAMs treated with miR-374b exosomes when compared to the corresponding control. Altogether, these results clearly suggest that regulation of EndoMT in SARSCoV-2 viral protein stimulated ECs is dependent on miR-374b (Fig. 5B and 5C).

## 4. Discussion

Corona virus disease (COVID-19) caused by SARS-CoV-2 acquired pandemic status making terrible impact on the public health. The fight against COVID-19 caused by SARS-CoV-2 infection is still raging. However, the pathophysiology of acute and post-acute complications of COVID-19 is still unclear [17]. Though initially recognized as an ARDS, COVID-19 is reported to be associated with high incidence of thrombosis and endothelial dysfunction [18]. Eventhough endothelial cell dysfunction is an important feature of severe COVID-19, the mechanisms through which SARS-CoV-2 induces endotheliopathy leading to thrombo-inflammatory and fibrotic damage still remain elusive. SARS-CoV-2 infection alters the balance of endothelial protective molecules and endothelial damaging molecules, leading to endothelial dysfunction[19]. SARS-CoV-2 mediated endothelial dysfunction can modify endothelial cell phenotype and contribute to endothelial-to-mesenchymal transition. Mounting evidences suggests endothelial to mesenchymal transition (Endo MT) is involved in fibrosis of COVID 19 patients in which endothelial cells (ECs) lose their endothelial specific properties and acquire mesenchymal properties, utimately leading to a pro-thrombotic state and endothelial dysfunction[20]. EndMT is regulated by complex molecular mechanisms including microRNAs (miRNAs)[21].

MicroRNAs (miRNAs), small endogenous RNA molecules that regulate several physiological processes including endothelial homeostasis, and vascular diseases, can be perturbed by infecting viruses. It has been reported that, SARS-CoV-2 infection can alter host miRNA expression landscape, resulting in miRNA mediated regulation of target mRNA and its downstream signaling cascade [22]. Eventhough host miRNAs are reported to be associated with SARS-CoV-2 infection, role of host miRNAs in regulating SARS-CoV-2 mediated endothelial dysfunction still remains elusive. Based on these facts, this study is designed to decipher the role of miRNAs in SARS-CoV-2 viral protein stimulated endothelial cells. In particular, this study focus on the role of miR-374b, which was found to be significantly downregulated upon profiling of SARS-CoV-2 viral protein, stimulated endothelial cells. Multiple studies have reported the alteration in host miRNA landscape following SARSCoV2 infection[23][24][25][26][27]. miR-374b is reported to be downregulated in blood samples of COVID-19 patients[16]. In order to identify the role of miR-374b in SARS-CoV-2 mediated endothelial dysfunction, the targets of miR-374b was predicted using online web based tools like miRDB, targetscan and miRsystem, Among the predicted targets, CFLAR, ITGB1 and VEGFA, only CFLAR could satisfy our criteria with a score of 3/3 and also CFLAR was found to be the most significantly upregulated gene in gene profiling, and hence was selected for further mechanistic studies. **CFLAR**, also known as c-FLIP, is a regulator of apoptosis that inhibits assembly of the caspase-8 and FADD complex. Cellular FLICE-inhibitory protein (c-FLIP) is a catalytically inactive caspase-8 (CASP8) homologue, which negatively regulates with apoptotic signaling [28]. Further, 3′UTR analysis results revealed that 3′UTR of c-FLIP contains two putative regions at nucleotides 6160–6167 and 10988–10994 that match the seed sequence of miR-374b, suggesting that 3′UTR of CFLAR possess potential binding sites for miR-374b-5p. CFLAR mRNA stability assay results showed that CFLAR mRNA stability was significantly reduced under miR-374b overexpression condition, suggesting that the regulatory role of miR-374b involves alterations in the post-transcriptional stability of CFLAR mRNA.

CFLAR(c-FLIP) being predicted as the target of miR-374b, and CFLAR mRNA and protein levels were increased in SARSCoV-2 viral protein stimulated ECs, we further checked the effect of miR-374b overexpression on CFLAR mRNA and protein levels in SARSCoV-2 viral protein stimulated ECs. Results of which showed that CFLAR mRNA and protein levels were significantly reduced in miR-374b overexpressing SARSCoV-2 viral protein stimulated ECs. CFLAR, being an anti-apoptotic protein, we further checked the effect of miR-374b on CFLAR mediated anti-apoptotic effect in SARSCoV-2 viral protein stimulated ECs. The results obtained suggest that ectopic expression of miR-374b increased the percentage of cells undergoing apoptosis in SARSCoV-2 viral protein stimulated ECs by reducing CFLAR levels. Since it was observed that, endothelial cell specifc markers like PECAM1, VWF and VE-CADHERIN was reduced in SARSCoV-2 viral protein stimulated ECs and that CFLAR is directly proportional to survival rate of cells, we next checked, if CFLAR can modulate the expression pattern of genes related to endothelial function/dysfunction, particularly genes associated with EndoMT like α SMA, Vimentin, FSP, and N-cadherin. Results of which showed that EndoMT markers were several fold increased in SARSCoV-2 viral protein stimulated ECs when compared to control cells. Further in order to check the effect of miR-374b mediated alteration of EndoMT markers, we next analysed the expression pattern of EndoMT markers in miR-374b overexpressing SARSCoV-2 viral protein stimulated ECs. Results obtained showed that EndoMT markers were significantly reduced in miR-374b overexpressing SARSCoV-2 viral protein stimulated ECs, suggesting that ectopic expression of miR-374b can reverse the expression pattern of EndoMT markers. In order to further prove the role of miR-374b in regulating Endo MT, chick Chorioallantoic membrane (CAM) assay was performed. CAM assay results showed that the expression level of Endo MT markers (SMA and FSP) increased in CAMs treated with SARSCoV2 spike protein was found to be reversed in CAMs co-treated with miR-374b exosomes. The micro vessel density and hemoglobin levels were found to be significantly lower in CAMs treated with miR-374b exosomes when compared to the corresponding control.

MiR-374b was found to be downregulated in blood samples of COVID-19 patients and our model system also mimicked the same. Further our study showed that, decreased levels of miR-374b and its regulation of c-FLIP through increased endothelial cell survival, it drastically affects the EC function as evidenced by induction of endoMT and this phenomenon may be responsible atleast in part for the endothelial dysfunction associated with SARSCoV2 infections. Altogether, these results clearly suggest that regulation of EndoMT in SARSCoV-2 viral protein stimulated ECs is dependent on miR-374b. Contrary to our observations and the available data from the clinical data on SARSCoV2, miR-374b has also been reported to induce endothelial to mesenchymal transition by inhibiting endothelial MAPK7 signalling in response to TGFβ [29]. This contradiction may be attributed to the context dependent specific regulation by miRNAs, and that the MAPK7 mediated regulation occurs in a different biological scenario associated with TGFβ signaling unlike the SARSCoV2 protein stimulation.

## 5. Conclusion

In conclusion, the data presented in this manuscript show that, miR-374b induces EndoMT in SARSCoV-2 viral protein stimulated ECs and also enhance cell survival by regulation of c-FLIP (Fig. 6).

**Fig. 6.**
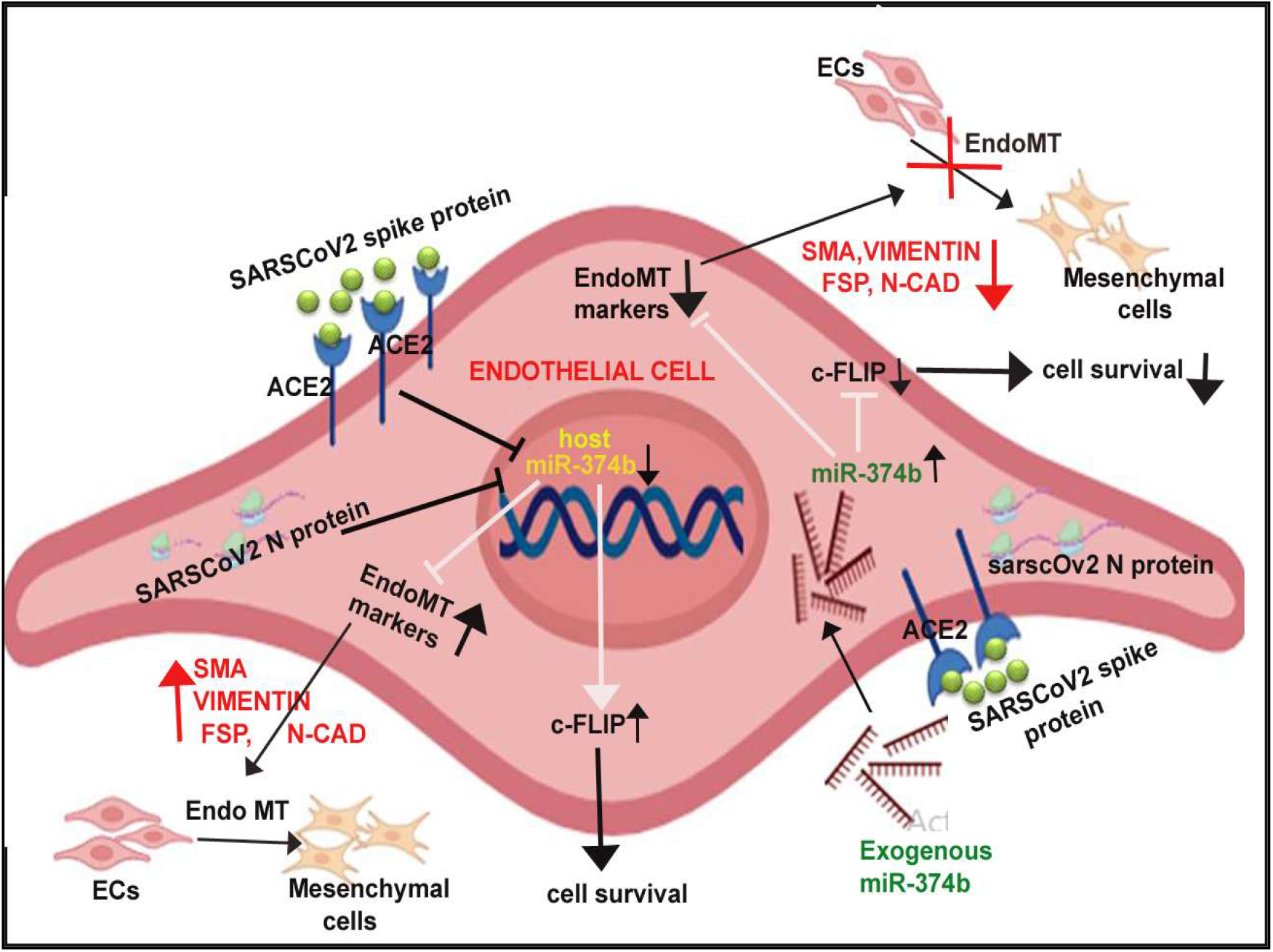
miR-374b dependent regulation of EndoMT in SARSCoV-2 viral protein stimulated ECs by targeting c-FLIP.

## Statements and Declarations

### Acknowledgement

Financial assistance received from Indian Council of Medical Research (ICMR), Department of Health Research, Govt. of India (Grant no. No. R.12013/23/2021-HR/E-Office: 8131577) is gratefully acknowledged.

## Author contributions

GRR did most of the experiments, helped in analysing the results and writing the manuscript. AP cloned miR-374b. AK and VR helped in CAM assay and data analysis. VBSK conceived the work, helped in analysing the results and writing the manuscript.

